## Supplementary information for "Cardiac stress leads to regulation of Filamin C dimerisation via an ancient phosphorylation-modulated interaction with HSPB7"

Extended methods

Extended Data Figs 1-8

Extended Data Tables 1-3

Extended references

#### Extended Methods

##### *Fitting titration data to extract dissociation constants and Gibbs free energies*

The titration data for the homo-dimerisation of the various $FLNC_{d24}$ constructs was collected by varying the total concentration of $FLNC_{d24}$ and measuring the ratio of the concentration of dimers to the total concentration. For the hetero-dimerisation of $FLNC_{d24}$ with the different HSPB7 constructs, the total concentration of FLNC was kept constant and titrated with varying concentrations of HSPB7. The abundances of the different species were extracted from the according peak heights in the mass spectra, and the ratio of the concentration of heterodimers to the total concentration of HSPB7 was calculated. The equilibrium constants for the dimerisation reactions were determined by fitting the experimental data to the simplest model for association as follows.

We defined two coupled equilibria for homo- and hetero-dimerisation with associated dissociation constants:

$$FLNC_2 \rightleftharpoons 2\,FLNC, \text{ where } K_{D,homo} = \frac{[FLNC]^2}{[FLNC_2]} \text{ (for homo-dimerisation)}$$

and

$$FLNC\colon HSPB7 \rightleftharpoons FLNC + HSPB7, \text{ where } K_{D,hetero} = \frac{[FLNC][HPSB7]}{[FLNC\colon HSPB7]} \text{ (for hetero-dimerisation).}$$

From the law of mass action, we know that:

$$[FLNC_{total}] = [FLNC] + 2[FLNC_2] \text{ (for homo-dimerisation)}$$

and

$$[FLNC_{total}] = [FLNC] + 2[FLNC_2] + [FLNC\colon HSPB7] \text{ and}$$
$$[HSPB7_{total}] = [FLNC\colon HSPB7] + [HSPB7]$$
$$\text{(for hetero-dimerisation).}$$

While the expression for the homo-dimerisation data can be solved exactly, that for hetero-dimerisation leads to a cubic equation with non-trivial solutions. We thus decided to solve both numerically using the Scipy *curve_fit* function which uses a non-linear least square fitting protocol.

First, we found $K_{D,homo}$ using the *fsolve* function from the *scipy.optimize* module that was used to find the equilibrium concentrations of FLNC monomers and homo-dimers that satisfy these equations for a given value of $K_{D,homo}$ and $[FLNC_{total}]$. The mole fraction of homo-dimers was then calculated, and the optimal value of $K_{D,homo}$ found by fitting the calculated mentioned ratio to the experimental data. This was then used as input for finding $K_{D,hetero}$. Upon knowing $[FLNC_{total}]$, $[HSPB7_{total}]$, $K_{D,homo}$, and $K_{D,hetero}$ the *fsolve* function was used to find the concentrations of $FLNC, FLNC_2, HSPB7$ and $FLNC\colon HSPB7$ that satisfy the law of mass action equations given above. The optimal value of $K_{D,hetero}$ was found by fitting the calculated fraction of HSPB7 in the heterodimer, $[FLNC\colon HSPB7]/[HSPB7_{total}]$ to the experimental data.

The fitted $K_Ds$ were used to calculate standard Gibbs free energies using $\Delta G^{\ominus} = -RTlnK_D$, where $R$ is the gas constant, 8.314 $Jmol^{-1}K^{-1}$ and $T$ the temperature, 298.15 K. Errors in as $\Delta G^{\ominus}$ were calculated using error propagation from the errors in fitting the titrations performed to obtain the respective $K_D$ values.

### Extended Data Figures

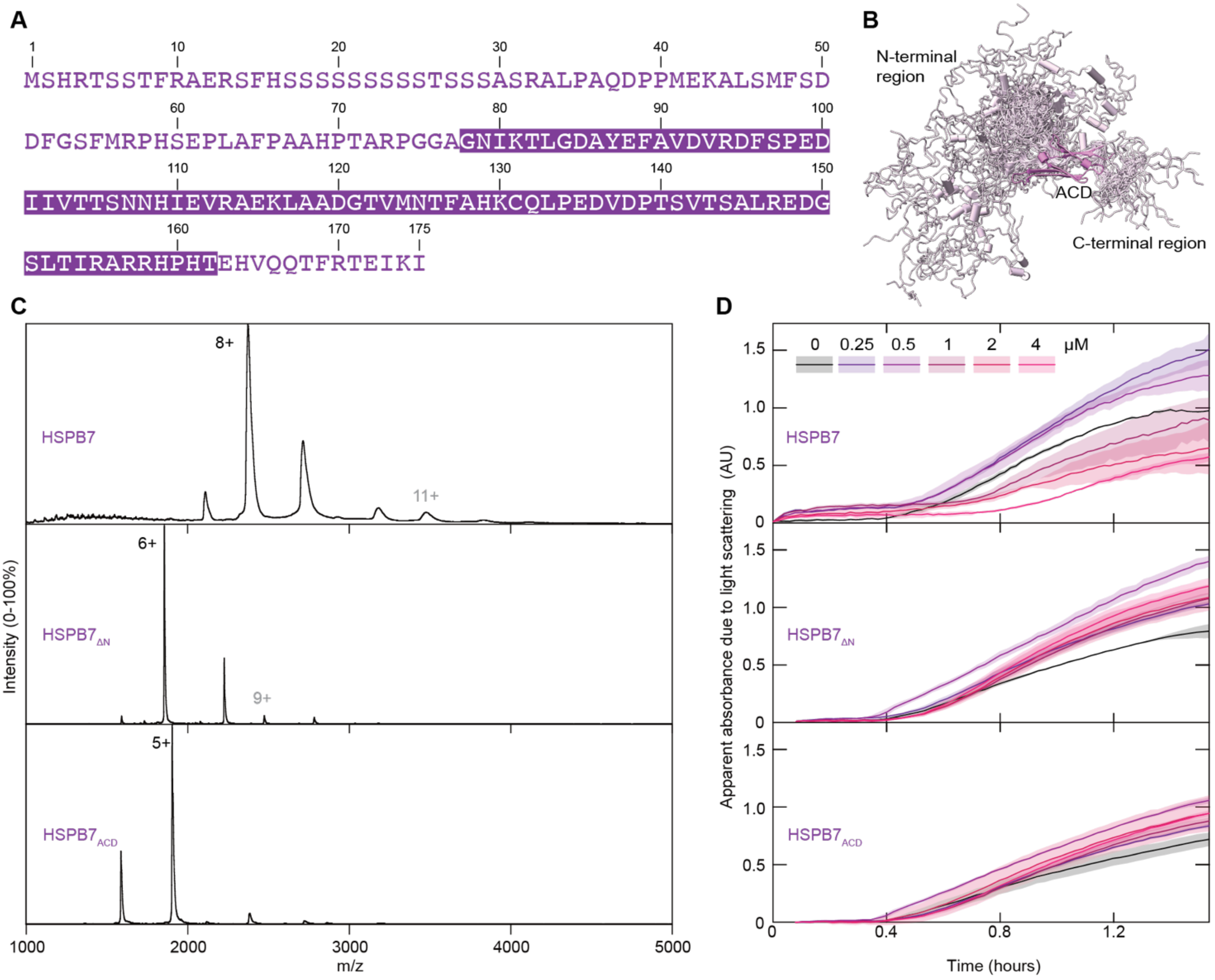


**Extended Data Fig. 1 | HSPB7 is predominately monomeric with weak chaperone activity. A** The sequence of HSPB7 (Uniprot ref: Q9UBY9-2, Isoform 2, A67 to AAHPTA, 175 residues), with the sequence for the ACD construct used in this work highlighted. **B** An ensemble of computational predictions using AlphaFold for the full-length HSPB7 structure, aligned to the structure of HSPB7$_{ACD}$, show the N- and C-terminal regions are likely disordered and heterogeneous. **C** Native mass spectra of the HSPB7 constructs at 10 µM show monomers (black charge state label) and only extremely low abundances of dimer (grey charge state) in all cases. **D** Chaperone assays in which the ability of HSPB7, HSPB7$_{\Delta N}$ and HSPB7$_{ACD}$ to attenuate heat-induced aggregation of citrate synthase (CS) was tested. The black curve reports the apparent absorbance due to light scattering due to the aggregation of 1 µM CS (without the addition of any of the HSPB7 constructs). Addition of HSPB7$_{\Delta N}$ and HSPB7$_{ACD}$, at concentrations up to 4 µM (purple to pink curves), does not decrease the aggregation of CS. Instead the apparent absorbance is somewhat increased, consistent with a weak interaction between the proteins leading to co-aggregation. For HSPB7, at the highest concentrations, the aggregation of CS was somewhat attenuated. These results show that the chaperone activity of HSPB7 is very limited, and relies heavily on the N-terminal region.

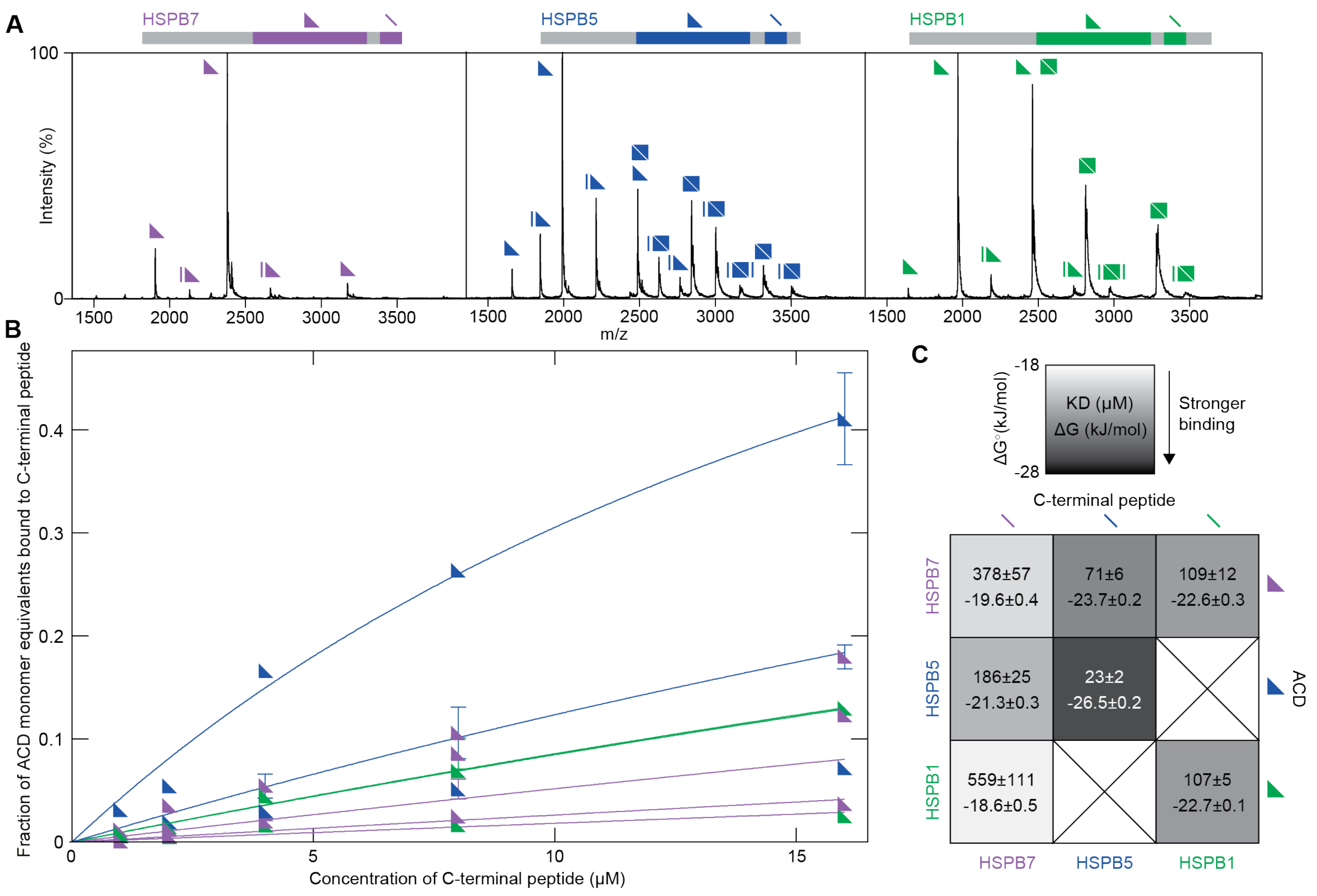


**Extended Data Fig. 2 | The HSPB7 C-terminal region does not bind to sHSP ACDs. A** Native mass spectra obtained when 5 µM of sHSP (HSPB7, purple; HSPB5, blue; HSPB1, green) ACD (triangle) were incubated with 40 µM of a peptide with their C-terminal sequence containing the IXI motif (line). The binding between $HSPB7_{ACD}$ and its own C-terminal peptide is very weak, compared with the same interaction in HSPB1 and HSPB5. **B** The titration experiments of the sHSP ACD-peptide interactions, the protein concentrations were maintained at 5 µM. The colours of the triangular symbols and line represent, respectively, the ACD peptide used in the titration (HSPB7, purple; HSPB5, blue; HSPB1, green). **C** The extracted thermodynamic parameters of sHSP ACD-peptide interactions. The matrix is shaded according to the strength of binding, and shows that the root cause of HSPB7 not binding its C-terminal peptide to a significant extent lies in the C-terminal, not in the ACD, sequence.

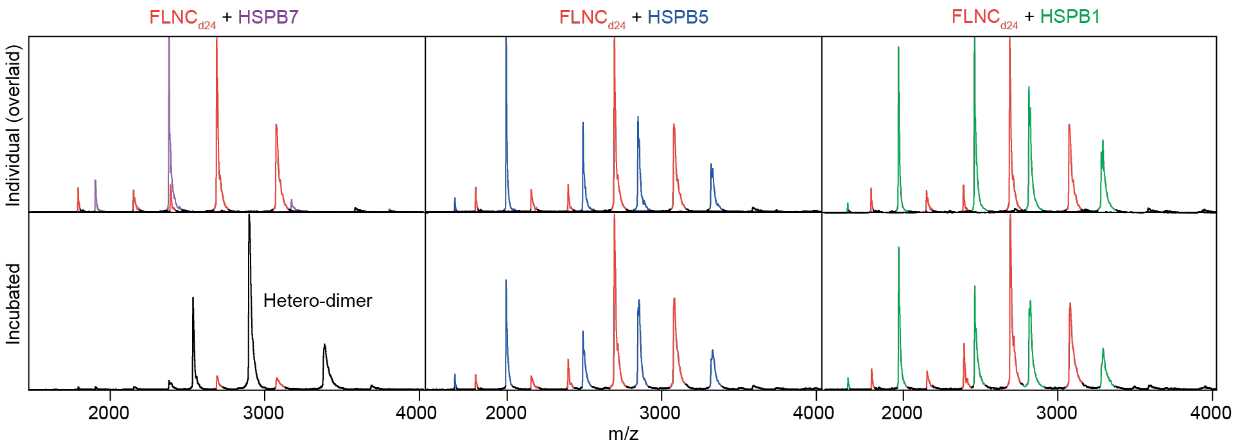


**Extended Data Fig. 3 | FLNC$_{d24}$ binds HSPB7$_{ACD}$ specifically. A** Native MS data of FLNC$_{d24}$ (red) incubated with HSPB7$_{ACD}$ (purple), HSPB5$_{ACD}$ (blue) or HSPB1$_{ACD}$ (green). The upper row shows the spectra of the individual proteins overlaid, the bottom row shows the interactions between proteins mixed at the monomer ratio of 1:1. Only HSPB7 forms observable amounts of hetero-dimer (black) with FLNC$_{d24}$. Experiments were performed under reducing conditions, given the ability of HSPB1 to form a covalently linked ACD homo-dimer.

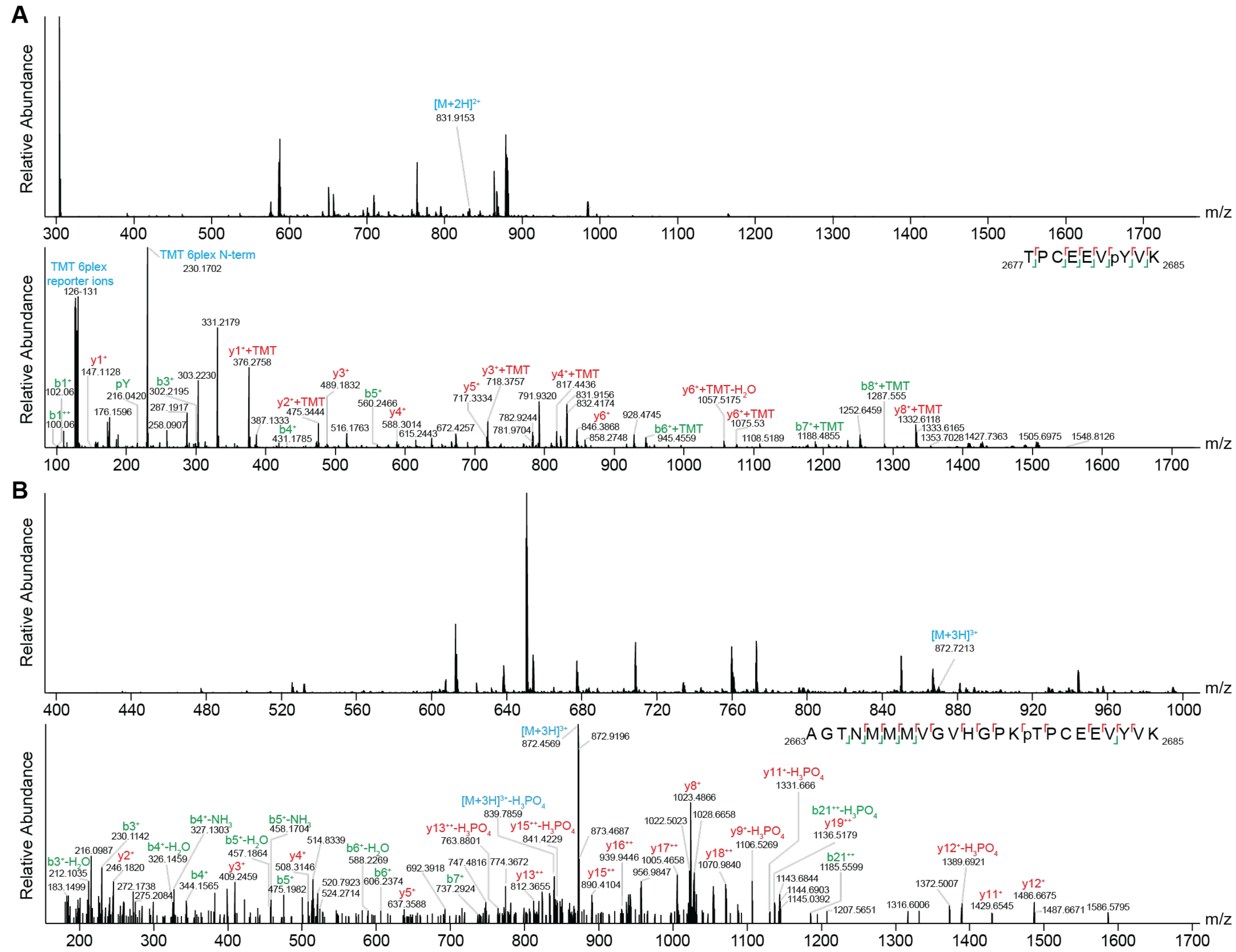


**Extended Data Fig. 4 | Annotated mass spectra of tryptic phosphopeptides of FLNC validate phosphorylation at T2677 and Y2683. A** MS1 mass spectrum (upper panel) with precursor peptide containing pY2683 selected for tandem-MS analysis indicated. The resulting MS2 mass spectrum (lower panel) displays many fragment ions, with those assignable to *b*- (green) and *y*-ions (red) annotated. The ion series provide phosphosite-specific information for the unambiguous localisation of pY2683. Data was extracted from the PRIDE database, having been deposited by Hoffmann et al[1]. TMT 6plex labeling was used in the study, hence TMT reporter ions and uncleaved TMT labels are also labelled. **B** MS1 mass spectrum (upper panel) with precursor peptide containing pT2677 selected for tandem-MS analysis indicated. The resulting "MS2" mass spectrum (lower panel) displays many fragment ions, with those assignable to *b*- (green) and *y*-ions (red) annotated. Fragment ions with a neutral loss of phosphoric acid (-$H_3PO_4$), ammonia (-$NH_3$), or water (-$H_2O$) are indicated in the MS2 spectra. The ion series provide phosphosite-specific information for the unambiguous localisation of pY2677. Data was extracted from the PRIDE database, having been deposited by Bouhaddou et al[2].

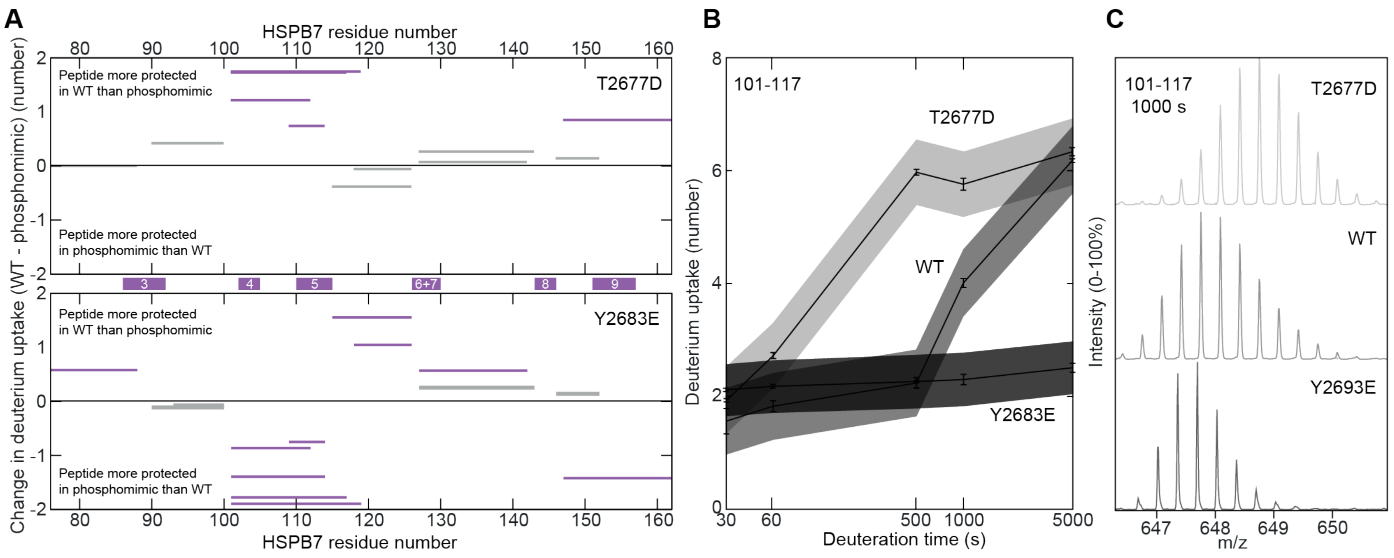


**Extended Data Fig. 5 | HDX reveals differences in solvent accessibility upon mimicking phosphorylation at the hetero-dimer interface. A** The Woods plots showing the differential deuterium uptake (after 1000 s of labelling) of $HSPB7_{ACD}$ hetero-dimerised with WT $FLNC_{d24}$, or one of the two phosphomimics T2677D (upper panel) and Y2683E. **B** A plot of deuterium uptake against deuteration time of the most protected peptide on $HSPB7_{ACD}$ (residues 101-117) in the different hetero-dimers. T2677D is the least protected, and Y2683E the most protected. This is in agreement with our native MS data that showed a weakening of the hetero-dimer interface in the former, and a strengthening in the latter, compared to the WT (**Fig. 4**). Error bars refer to the standard deviation of three repeats at each time-point, and the shaded bands a 99% confidence band. **C** Raw mass spectra of the $HSPB7_{ACD}$ peptide (residues 101-117), in the 3+ charge state at 1000 s, for the each of the three complexes shows how the differences in deuteration are readily visible by eye.

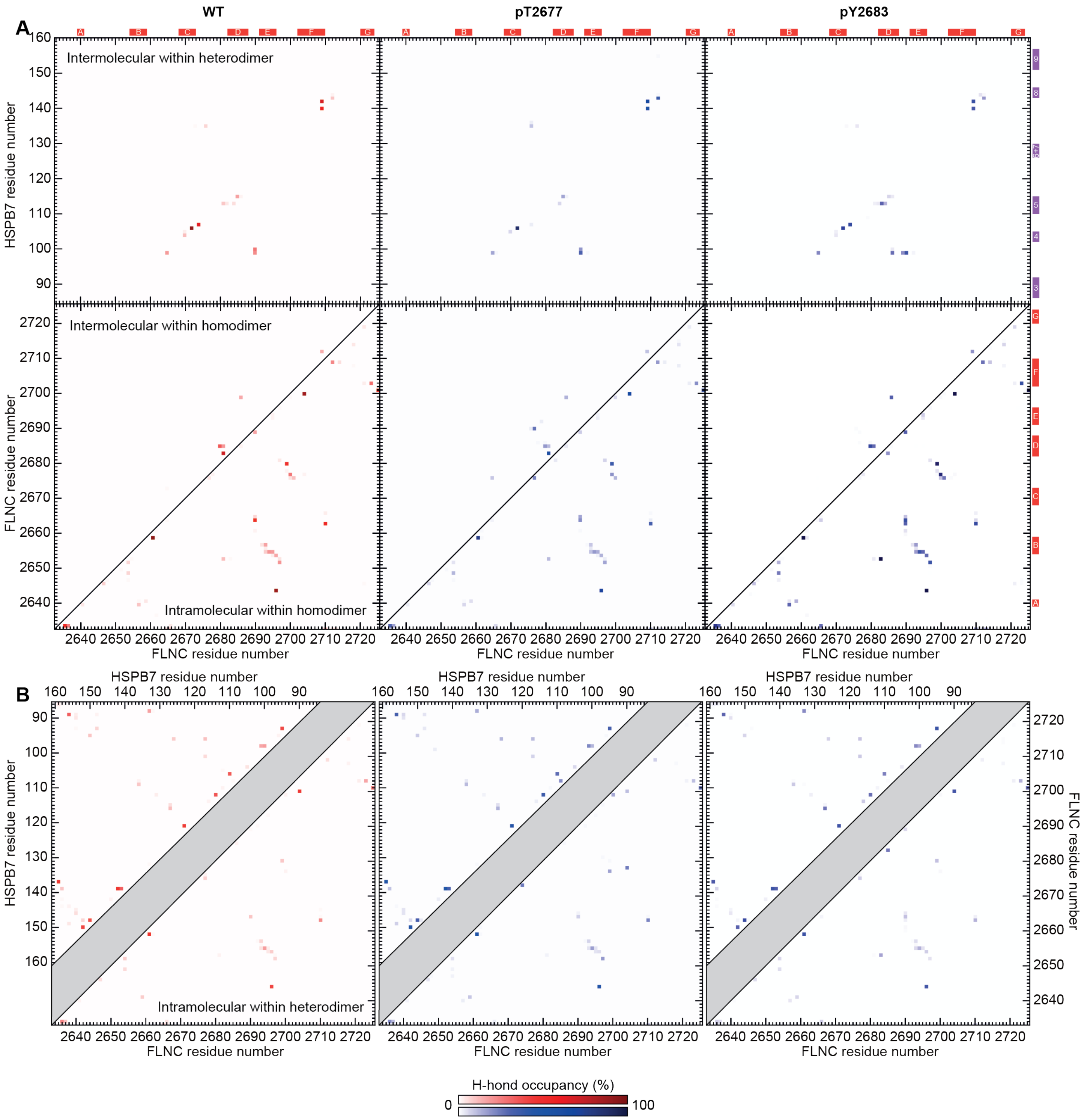


**Extended Data Fig. 6 | Hydrogen-bond maps for WT and phosphorylated FLNC$_{d24}$ homo- and hetero-dimers.** The data underlying the difference maps shown in **Fig. 5** and **Extended Data Fig. 8**. **A** Bond occupancy for inter-molecular hydrogen bonds within the hetero-dimer (upper row), and intra-molecular within the homo-dimer (lower row), for WT, pT2677 and pY2683 (from left to right). The darker the pixel, the larger the percentage of frames within the MD simulation in which a hydrogen bond is formed between those two residues. **B** Bond occupancy for intra-molecular hydrogen bonds within the hetero-dimer. Bonds within HSPB7 are shown in the upper left triangles, and bonds within FLNC are shown in the lower right triangles. Data for WT, pT2677 and pY2683 (from left to right) are displayed using the same colour scale as for **A** (shown at the bottom).

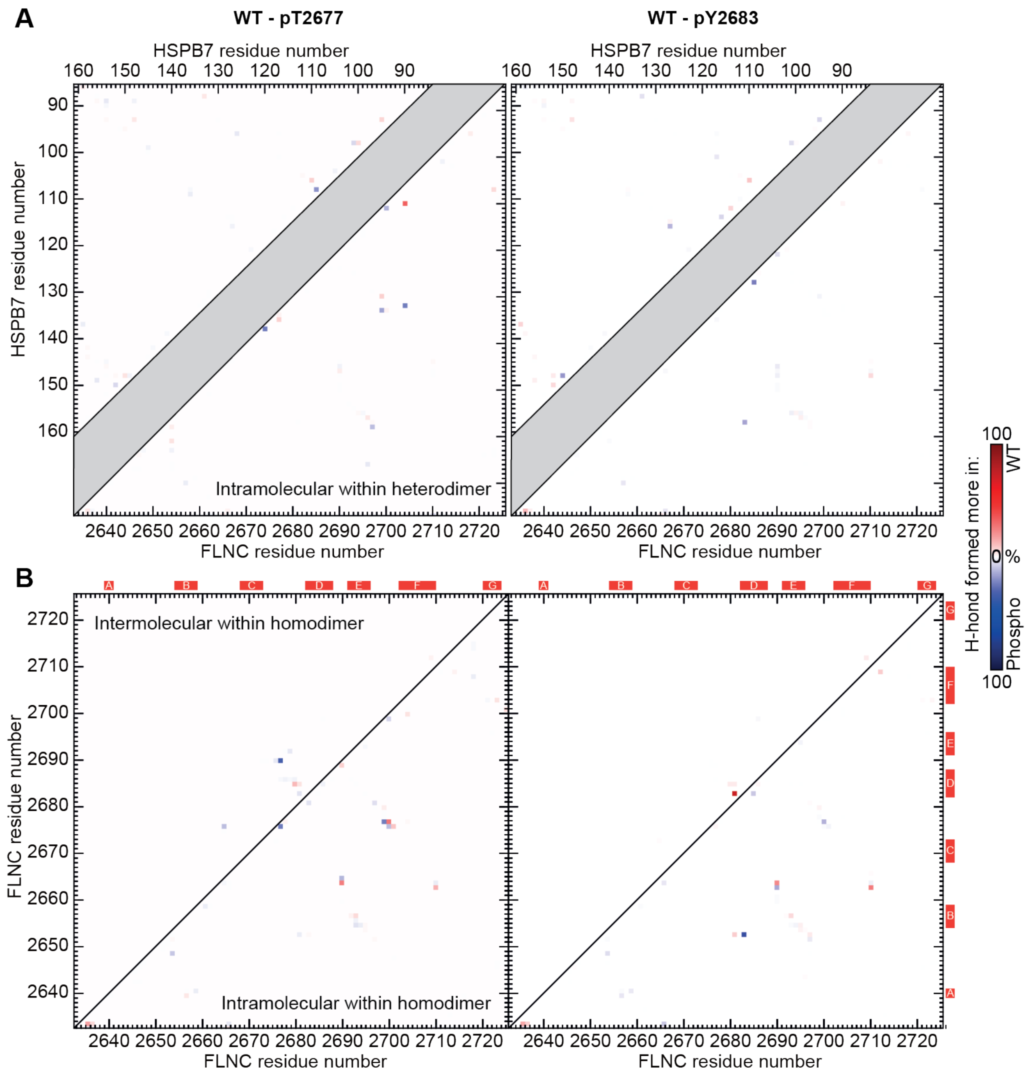


**Extended Data Fig. 7 | Hydrogen-bond difference maps for WT and phosphorylated FLNC$_{d24}$ homo- and hetero-dimers. A** Difference in bond occupancy for intra-molecular hydrogen bonds within the hetero-dimer between WT and pT2677 (left) or pY2683 (right). Bonds within HSPB7 are shown in the upper left triangles, and bonds within FLNC are shown in the lower right triangles. **B** Difference in bond occupancy for inter-molecular (upper left triangles) and intra-molecular (lower right triangles) hydrogen bonds within the homo-dimer between WT and pT2677 (left) or pY2683 (right). Colour scale as for **A** (shown to the right).

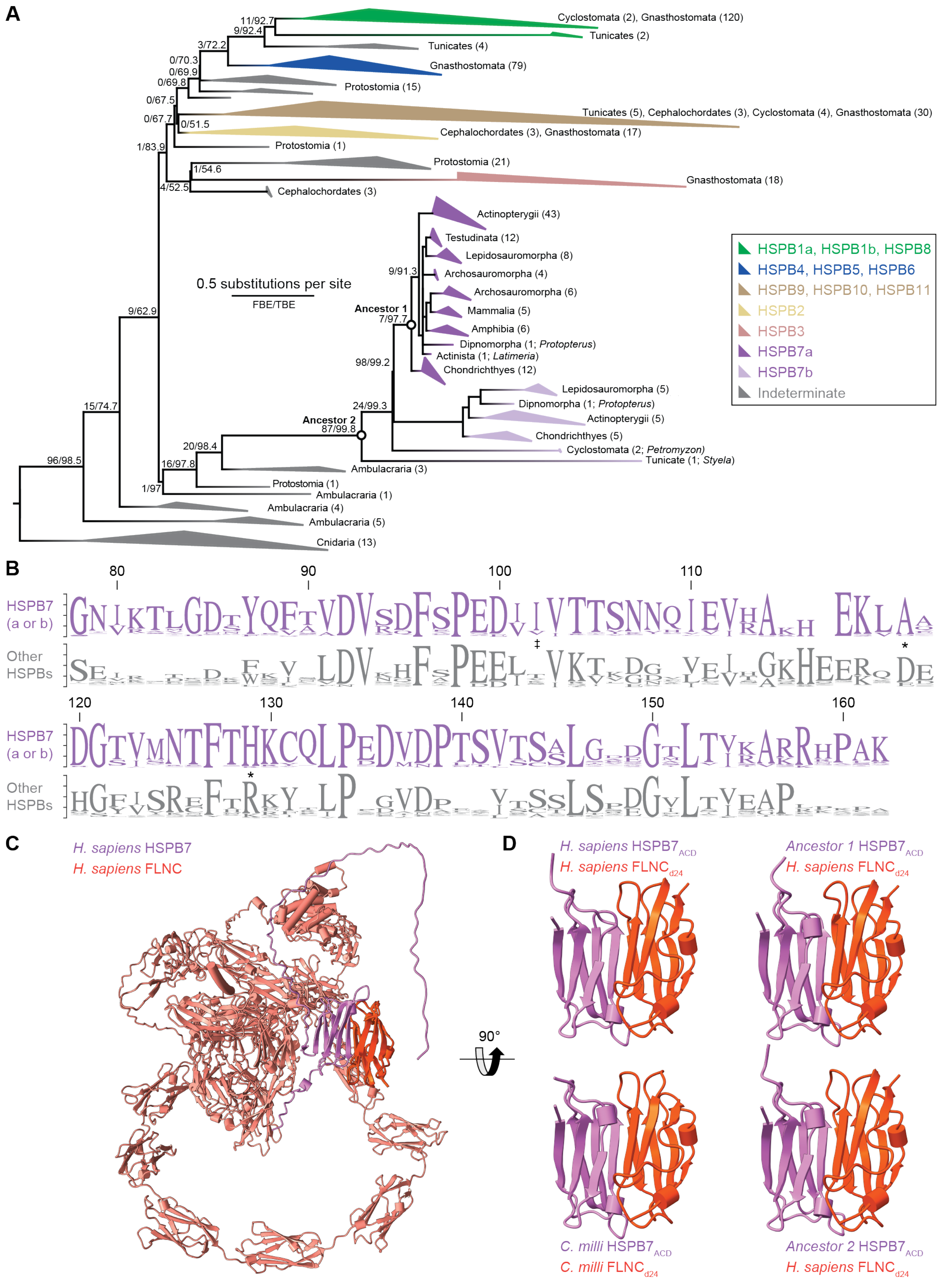


**Extended Data Fig. 8 | The interaction between FLNC and HSPB7 is ancient. A** Maximum likelihood phylogeny of HSPBs based on searching for all of HSPB1-11. HSPB7 can be identified in species as distantly related to humans as tunicate *Styela*, and is present in all vertebrate clades in between. We find two distinct forms of HSPB7, which we term HSPB7a and HSPB7b, with humans having only the first type. Ancestral proteins that we reconstructed are indicated at their corresponding nodes on the tree. **B** Sequence logo of the frequency of different amino acids in the ACD for all the HSPB7s and other HSPBs (grey) in our phylogeny. The two sites we identified (**Fig. 2C**) as likely responsible for monomerisation are marked (*), as is the site of I102 (in human HSPB7) that is involved in the hydrophobic interface of the heterodimer (‡). **C** AlphaFold model of a single chain of full-length human FLNC (orange; d24 in a darker hue) in complex with a single chain of HSPB7 (purple). The interface we observed in our structure of $FLNC_{d24}$:$HSPB7_{ACD}$ (**Fig. 3B**) is preserved in this model, demonstrating that the missing domains of FLNC and terminal regions of HSPB7 do not preclude hetero-dimerisation. **D** AlphaFold models of the $FLNC_{d24}$:HSPB7 hetero-dimer for the human and *C. milli* proteins (left), and the human $FLNC_{d24}$ with ancestral forms of HSPB7 (right) in our competitive binding assay. Only one chain of each protein is shown in these models, and the terminal regions of HSPB7 are hidden, for clarity. All four cases form hetero-dimers that align essentially perfectly with our crystal structure (**Fig. 3B**).

#### Extended Data Tables

| | $HSPB7_{ACD}^{C131S}$ | $FLNC_{d24}$:$HSPB7_{ACD}^{C131S}$ |
|---|---|---|
| **PDB ID** | 8RHA | 8PA0 |
| **Data collection** | | |
| Space group | $P4_12_12$ | $P3_221$ |
| Cell dimensions | | |
| *a*, *b*, *c* (Å) | 57.86, 57.86, 167.34 | 94.13, 94.13, 51.60 |
| α, β, γ (°) | 90, 90, 90 | 90, 90, 90 |
| Synchrotron beamline | DLS I04 | DLS I03 |
| Data collection temperature (K) | 100 | 100 |
| Wavelength (Å) | 0.97950 | 0.97950 |
| Resolution (Å) | 55.78-2.18 (2.22-2.18) * | 47.04-2.85 (2.90-2.85) * |
| No. total reflections | 398937 (19156) | 122811 (6265) |
| No. unique reflections | 15713 (764) | 6383 (316) |
| $R_{merge}$ | 0.138 (1.418) | 0.288 (2.487) |
| $I / \sigma I$ | 14.5 (2.2) | 6.1 (1.9) |
| Completeness (%) | 100 (99.1) | 100 (100) |
| Redundancy | 25.4 (25.1) | 19.2 (19.8) |
| CC1/2 | 1 (0.957) | 0.995 (0.654) |
| **Refinement** | | |
| Resolution (Å) | 54.68-2.18 (2.22-2.18) | 47.06-2.85 (2.90-2.85) |
| $R_{work}$ / $R_{free}$ | 0.2534/0.2796 | 0.2243/0.2909 |
| No. atoms | | |
| Protein | 1,807 | 1,350 |
| Ligand/ion | 0 | 0 |
| Water | 6 | 4 |
| Average *B*-factors <Å$^2$> | | |
| Protein | 62.78 | 59.93 |
| Ligand/ion | / | / |
| Waters | 46.06 | 51.37 |
| R.m.s. deviations from ideal | | |
| Bond lengths (Å) | 0.006 | 0.005 |
| Bond angles (°) | 1.399 | 0.54 |
| Ramachandran favored (%) | 97.33 | 94.19 |
| Ramachandran allowed (%) | 2.67 | 5.81 |
| Ramachandran outliers (%) | 0.00 | 0.00 |

*Values in parentheses are for highest-resolution shell.

**Extended Data Table 1 | Data collection and refinement statistics for $HSPB7_{ACD}^{C131S}$ and $FLNCd_{24}$:$HSPB7_{ACD}^{C131S}$.**

| | Theoretical (Da) | Experimental (Da) |
|---|---|---|
| FLNC$_{\text{d24}}$ monomer | 10747.4 | 10758.7±1.4 |
| FLNC$_{\text{d24}}$ dimer | 21494.8 | 21519.4±0.7 |
| FLNC$_{\text{d24}}^{\text{T2677D}}$ monomer | 10761.4 | 10775.5±0.7 |
| FLNC$_{\text{d24}}^{\text{T2677D}}$ dimer | 21522.8 | 21544.9±1.4 |
| FLNC$_{\text{d24}}^{\text{Y2683E}}$ monomer | 10713.4 | 10723.5±2.1 |
| FLNC$_{\text{d24}}^{\text{Y2683E}}$ dimer | 21426.7 | 21447.4±0.7 |
| HSPB7$_{\text{ACD}}$ | 9507.5 | 9517.2±1.2 |
| FLNC$_{\text{d24}}$:HSPB7$_{\text{ACD}}$ | 20254.9 | 20277.6±3.2 |
| FLNC$_{\text{d24}}^{\text{T2677D}}$:HSPB7$_{\text{ACD}}$ | 20268.9 | 20294.4±2.1 |
| FLNC$_{\text{d24}}^{\text{Y2683}}$:HSPB7$_{\text{ACD}}^{\text{E}}$ | 20220.9 | 20241.9±2.8 |

**Extended Data Table 2 | Theoretical and experimental masses of the complexes.** Theoretical masses are calculated directly from the sequence. Experimental masses were obtained from native MS, and are measured from the peak centres. Native MS data were obtained under soft desolvation conditions so as not to artefactually dissociate the complexes, which leads to residual binding of buffer components. Consequently somewhat higher masses (typically <1%[3]) are measured than predicted by sequence alone. However, these mass shifts are small relative to the mass differences between our assignments and any other alternatives given the few components in each mixture.

| | HSPB7$_{\text{ACD}}$ | FLNC$_{\text{d24}}$ | Distance (Å) |
|---|---|---|---|
| **Salt bridges** | D100 OD1 | H2686 ND1 | 2.64 |
| | D100 OD2 | H2686 ND1 | 3.86 |
| **Hydrogen bonding** | E99 O | Y2692 HH | 1.78 |
| | V103 H | M2668 O | 2.57 |
| | V103 O | V2670 H | 2.38 |
| | T105 H | V2670 O | 2.41 |
| | T105 O | V2672 H | 2.48 |
| | N107 HD21 | G2674 O | 1.64 |
| | R113 HH21 | V2684 O | 1.87 |

**Extended Data Table 3 | Residue pairs forming hydrogen bonds and salt bridges across the interface in the FLNC$_{\text{d24}}$:HSPB7$_{\text{ACD}}^{\text{C131S}}$ crystal structure.**

#### Extended References


1. Hoffman, N. J. *et al.* Global Phosphoproteomic Analysis of Human Skeletal Muscle Reveals a Network of Exercise-Regulated Kinases and AMPK Substrates. *Cell Metab* **22**, 922–935 (2015).
2. Bouhaddou, M. *et al.* The Global Phosphorylation Landscape of SARS-CoV-2 Infection. *Cell* **182**, 685-712.e19 (2020).
3. Benesch, J. L. P. & Ruotolo, B. T. Mass spectrometry: Come of age for structural and dynamical biology. *Curr Opin Struc Biol* **21**, 641–649 (2011).
